# Sex-dependent gene regulation of human atherosclerotic plaques by DNA methylation and transcriptome integration points to smooth muscle cell involvement in women

**DOI:** 10.1101/2021.01.28.428414

**Authors:** Robin J. G. Hartman, Marten A. Siemelink, Saskia Haitjema, Koen F. Dekkers, Lotte Slenders, Arjan Boltjes, Michal Mokry, Nathalie Timmerman, Gert J. de Borst, Bastiaan T. Heijmans, Folkert W. Asselbergs, Gerard Pasterkamp, Sander W. van der Laan, Hester M. den Ruijter

## Abstract

Sex differences are evident in the clinical presentation and underlying histology of atherosclerotic disease with women developing more stable atherosclerotic lesions than men. It is unknown whether this is explained by sex differences in gene regulation in cellular compartments of atherosclerotic plaques. To study sex differences in gene regulation we performed genome-wide DNA methylation and transcriptomics analysis on plaques of 485 carotid endarterectomy patients (31% female). Sex-differential DNA methylation at 4,848 sites in the autosome was enriched for cell-fate commitment and developmental processes, and its deconvolution predicted more smooth muscle cells in females, as compared to more immune cells in males. RNA-sequencing of the same plaques corroborated the sex differences in DNA methylation predicted cell-types, in which genes that were higher expressed in females were enriched for TGF-beta signaling and extracellular matrix biology. In addition, female-biased genes were enriched for targeting by regulatory loci based on sex differential methylation. Lastly, by using single-cell RNA sequencing we showed that these female-biased genes are mostly expressed in smooth muscle cells, and higher expressed in smooth muscle cells from female (predominantly stable) plaques as compared to male (relatively unstable) plaques. Our approach identified female-biased genes in smooth muscle cells in fibrous atherosclerotic plaques. This points towards new mechanisms in smooth muscle cell biology of stable atherosclerotic plaques and offers new directions for research to develop new sex-specific therapeutics for atherosclerotic disease.

## Introduction

Men and women differ in predisposition to cardiovascular disease (CVD), as well as its pathophysiology, prognosis and outcomes^1^. Thrombus formation superimposed on atherosclerotic plaque complicates and accelerates luminal stenosis and often precedes CVD. The underlying lesion composition may vary, and differences in plaque characteristics have been consistently reported between men and women^2–5^. Histological analyses of carotid plaques of both sexes point to more stable lesions in women, coinciding with less inflammation, more smooth muscle cells (SMC), and better prognosis in women as compared to men^2–5^. These sex-associated differences in plaque characteristics coincide with a different mechanism of eliciting CVD, as men tend to develop more plaque rupture as a substrate for CVD, while women have been shown to suffer more from stable plaques and plaque erosion^6^. The mechanisms underlying these sex differences in atherosclerotic lesions are mostly unknown, and studies rarely take the effect of sex into account in the research domain of cardiovascular epigenetics^7^. We hypothesized that sex differences in gene regulation and transcription in part underlies atherosclerotic lesion differences between the sexes, pointing towards new cell- and sex-specific mechanisms of atherosclerotic disease.

Therefore, we investigated both autosomal DNA methylation (DNAm) and RNA expression of carotid artery plaques of 485 patients (31% female) undergoing carotid endarterectomy by sex. We explored if sex differences in DNAm are linked to sex differences in cell composition by deconvolution analysis. Next, we studied the sex-specific corresponding RNA expression within the same atherosclerotic plaques.

Sex-differences herein were investigated further at a single-cell level using sex-stratified single-cell RNA sequencing to uncover sex- and cell-specific pathways of atherosclerotic lesions.

## Materials and methods

### Study population

The Athero-Express Biobank (AE), of which the study design has been published before^8^ is an ongoing biobank study that includes patients undergoing arterial endarterectomy. For this study, patients were included who underwent a carotid endarterectomy and of whom genotyping data were available. All methods regarding sample collection, atherosclerotic plaque histology, DNA extraction and methods regarding the normalization of the DNA methylation data have been published previously^9^. We used one of the cohorts for our main analyses (AEMS450K1, Athero-Express Methylation Study Illumina 450K 1) and the other smaller cohort for replicating the effect size of methylation sex differences (AEMS450K2, Athero-Express Methylation Study Illumina 450K 2). RNA-sequencing was performed on plaques for which DNA methylation data was available (420 out of 485 plaques). Carotid plaques from 37 (26 male and 11 female) patients from the AE biobank were used for single-cell sequencing.

Patient data can be found in the Supplemental Dataset.

The performed study is in line with the Declaration of Helsinki and informed consent was provided by all study participants, after the approval for this study by medical ethical committees of the different hospitals was obtained.

### Computational analysis

Extensive quality control of the methylation data was performed as described previously^9^. We did not investigate sex differences on the sex chromosomes, as the inactive and heterochromatic X-chromosome in females is littered with DNAm, and the same holds true for the heterochromatic Y-chromosome in males. As these phenomena would lead to major sex differences complicating analysis, we removed the probes located on the sex chromosomes. CpGs that overlapped with SNPs (based on data from GoNL) and those with multiple mappings were removed^10^. Furthermore, we removed probes that previous studies have shown to be dubious in quality. In the end, we were left with 275,204 probes. We calculated differential methylation between the sexes (Sex-differentially methylated CpG-probes, abbreviated as sexDMPs) using the limma package, while correcting for age and hospital of inclusion. CpGs were annotated using ChIPseeker^11^, with a transcription start site ranging from −3,000 to 3,000 bp of the start of the gene. Coverage plots (Fig. 2) and feature plots (Fig. 2 and Fig. 3B) were generated using ChIPseeker^11^. Analyses were performed in R (version 3.5.1), plots were generated in R as well. To determine potential regulatory functions of the sexDMP loci, we used the T-Gene algorithm in combination with the ENCODE hg19 Tissue Panel data^12^. In short, T-Gene predicts which genes are regulated by genomic loci based on readily available histone modification and transcription data. It calculates a score combining distance and correlation between histone modification data based on ChIP-sequencing and RNA expression. The loci we used encompassed 50 basepairs centered around the sexDMPs in the genome.

**Fig 1.**
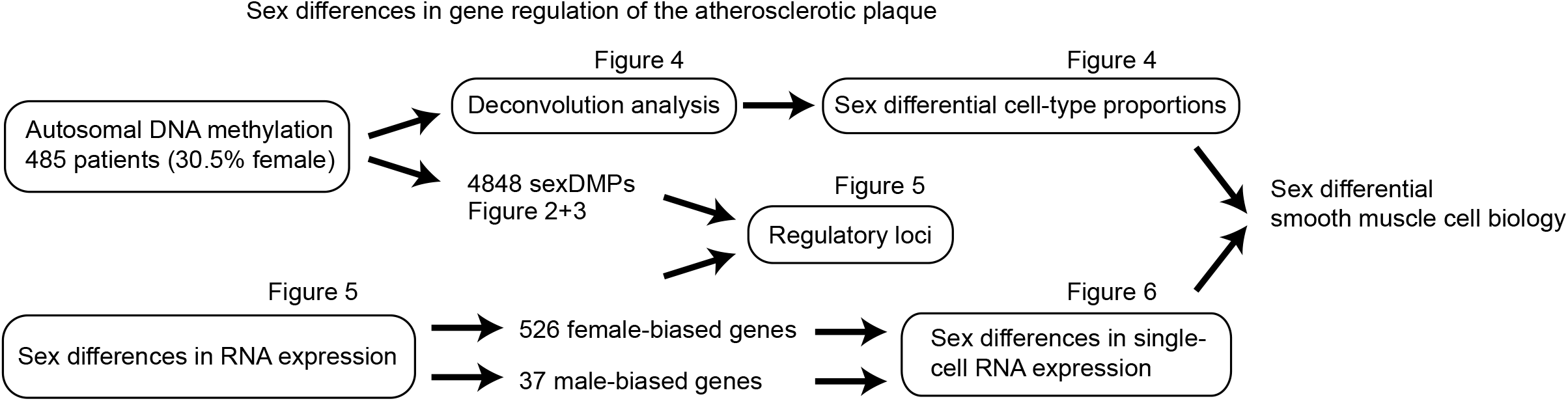
Flow-chart study. A flow chart is shown depicting the different analyses performed for this study.

**Fig 2.**
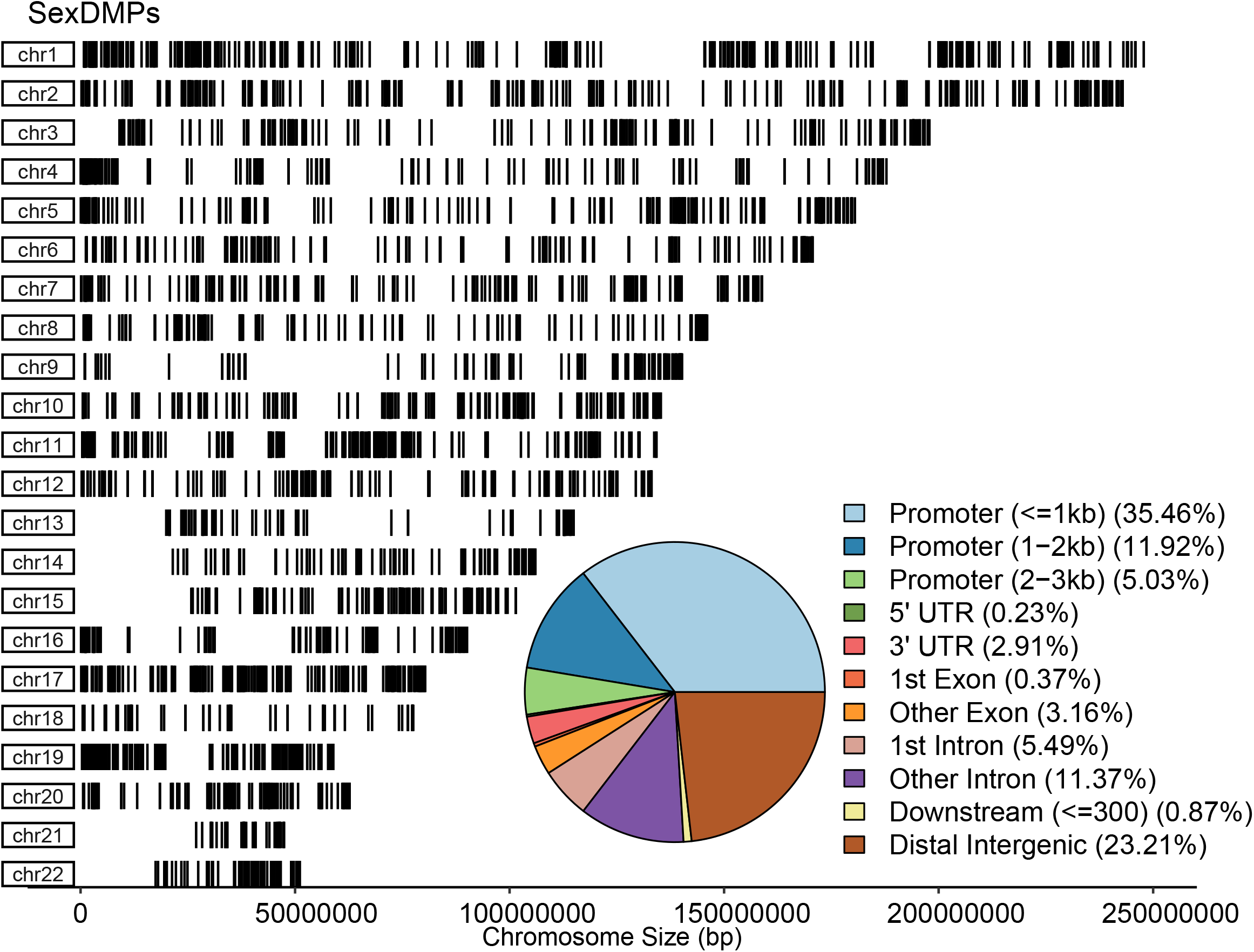
Sex differences in DNA methylation. A) A coverage plot of sex differentially methylated sites (sexDMP) over the autosome is shown. Each black line indicates one sexDMP. Chromosomal size in base pairs is shown on the X-axis, while the rows show the chromosome. The pie-chart shows the annotation of the location of the sexDMPs with respect to genes. The colored legend highlights the different annotation, e.g. lightblue = promoter (<1 kb).

**Fig 3.**
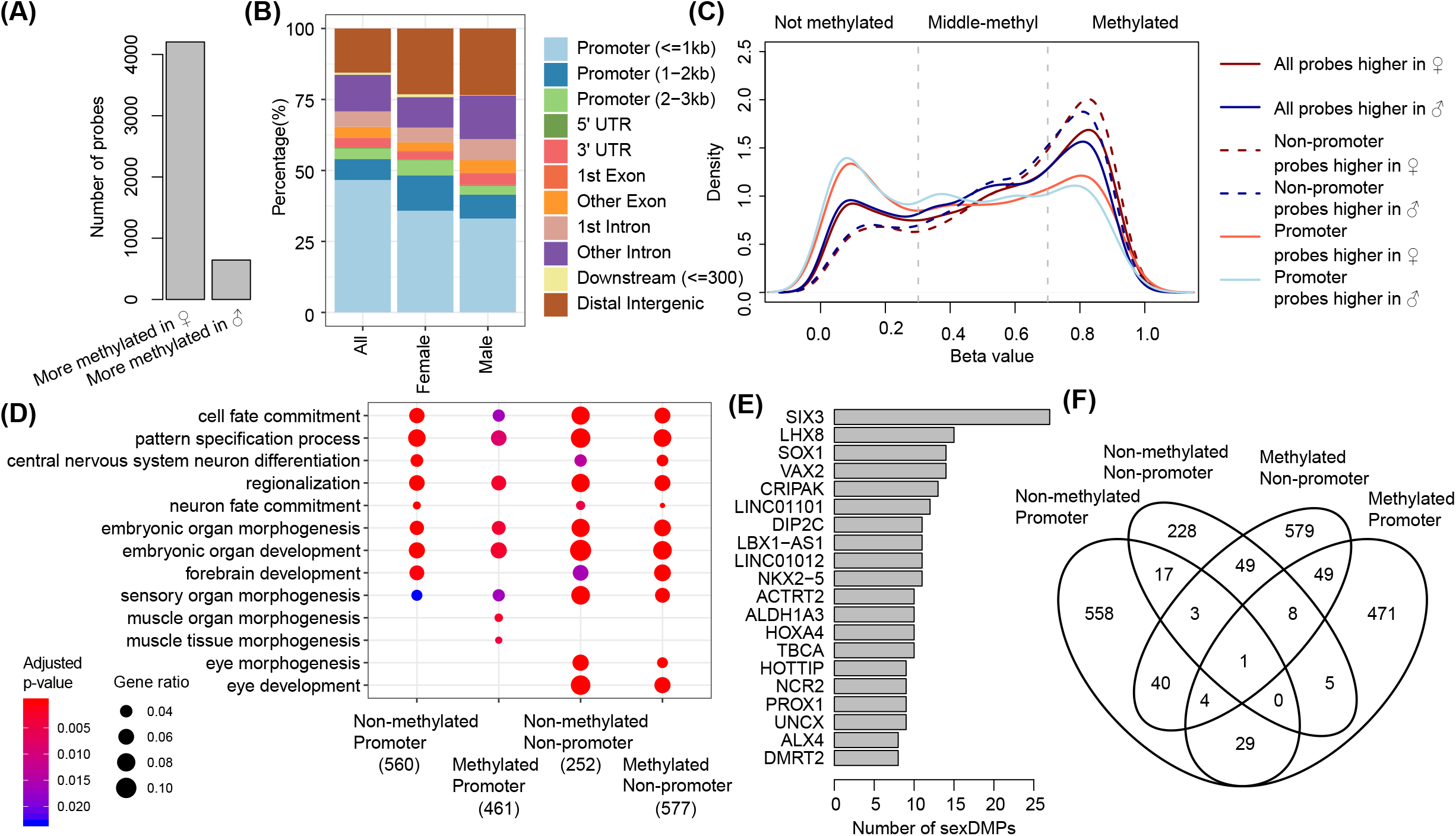
Direction of effect and underlying biology of sex differential methylation. A) The number of sexDMPs with either higher methylation values in females (left) or males (right) is shown in barplot. B) Barcharts show the annotation of the location of the sexDMPs with respect to genes split over direction of effect and for all CpGs tested. Colors indicate different annotations. From left to right the barcharts show all CpGs tested, those with higher methylation values in females, and lastly those with higher methylation values in males. C) The distribution of beta values of sexDMPs is shown in density plots. Blue and red lines indicate sexDMPs with higher methylation values in males and females, respectively. The darker uninterrupted lines show the distribution pattern for all sexDMPs, the lighter uninterrupted lines show the pattern for promoter sexDMPs, and the dashed lines show the pattern for non-promoter sexDMPs. Three different types of methylation states are separated by a vertical dashed line at beta-values of 0.3 and 0.7. D) A dotplot is shown for Gene Ontology enrichments in four different sexDMP sets separated by methylation state and whether or not the sexDMP is located in a promoter. Terms are allocated to the rows, color indicates significance, and size of the dot indicates the ratio of genes from the set present. The number indicates the number of genes that could be found in any of the sets tested. E) The top 20 genes and the amount of sexDMPs mapped to them are shown in a barplot. F) A venn-diagram is shown highlighting the overlap of genes mapped to by sexDMP sets separated by methylation state and whether or not the sexDMP is located in a promoter.

### Deconvolution analysis

Deconvolution of the DNA methylation data was performed using RefFreeEWAS^13^ with a k = 7, based on the number of main cell-types we expected in the atherosclerotic plaque (endothelial cells, SMCs, macrophages, mast cells, T cells, B cells, and a left-over “bin” cell-type). We performed a reference free deconvolution using the beta-values of the methylation data featuring 275,204 probes. Correlation plots of the μ (predicted cell-type specific methylation pattern) and Ω-matrices (predicted cell-type proportions per sample) were generated using the corrplot package. Gene enrichment analyses were performed using clusterProfiler^14^ for gene ontology biological processes. Gene enrichment analysis to determine cell-type for male and female-biased cell proportions were performed on CpGs limited to promoters that were non-methylated (< 0.3 beta-value) in female-biased versus methylated in male-biased cell-types (> 0.7) and vice versa. Sex-bias in cell-type proportions was determined by a statistically significant difference of p < 0.05 (Welch two-sample T-test) between the sexes.

### RNA-sequencing

Plaque segments were thawed, cut up, and further homogenized using ceramic beads and tissue homogenizer (Precellys, Bertin instruments, Montigny-le-Bretonneux, France), in the presence of TriPure (Sigma Aldrich), and RNA was isolated according to TriPure manufacturer’s protocol. From here, RNA in the aqueous phase was precipitated using isopropanol, and washed with 75% ethanol, and subsequently stored in 75% ethanol for later use or used immediately after an additional washing step with 75% ethanol. Next, libraries were prepared according to the CEL-Seq2 protocol^15^, with adaptations according to^16^. Libraries were sequenced as 2 x 75bp paired-end on a NextSeq 500 (Utrecht Sequencing Facility). The reads were demultiplexed and aligned to human cDNA reference using the BWA^17^ (0.7.13). Multiple reads mapping to the same gene with the same unique molecular identifier (UMI, 6bp long) were counted as a single read.

RNA-sequencing count data was corrected for UMI-count by rounding 4,096*(log(1-(gene counts / 4,096))). Next, only genes with a mean expression of higher than 1 were used for downstream analysis. We only used samples of which DNA methylation data was also present, leading to 420 samples. Outlying samples were removed based on hierarchical clustering of the distance matrix of the gene expression using cutreeStatic from the WGCNA package^18^ at a height of 17,500 after careful inspection of the dendrogram. This step removed 15 outlying samples. Sex differential gene expression was calculated using DESeq2^19^ while correcting for hospital of inclusion and age. A gene was called differentially expressed if *p* < 0.05. Gene enrichment analyses were performed using clusterProfiler^14^ for gene ontology biological processes. Backtracking female-biased genes to cell-type specific DNA methylation patterns was performed using only promoter CpGs within an absolute distance to the transcription start site of less than 1,000 bp. Next, the average of these promoter CpGs was taken of the μ-matrix of the deconvolution analysis.

### Single cell RNA sequencing data

Plaque samples minus the necrotic core from 11 females and 26 males were processed immediately after surgery, washed in RPMI and minced prior to digestion, as described in ^20,21^. The cell suspension was filtered through a 70 μm cell strainer and washed with RPMI 1640. Cells were kept in this medium with 1% Fetal Calf serum until staining for fluorescence-activated cell sorting. Single cell suspensions were stained with Calcein AM and Hoechst (ThermoFisher Scientific) in PBS supplemented with 5% Fetal Bovine Serum and 0.2% EDTA and subsequently strained through 70 μm cell strainers. Only cells positive for both Calcein AM and Hoechst (= living cells) were sorted for sequencing. Cells were prepped using the SORT-seq protocol^22^ and libraries were constructed using the CEL-seq2 protocol^15^. Data was processed in an R 3.5 environment using the Seurat package (v3.1.5)^23^. Downstream processing was performed using custom R scripts. The module score for female-biased genes was calculated using the addModuleScore function in Seurat. Sex differential module score in ACTA2+ SMCs was determined using a Welch two sample T-test in R. Sex differential gene expression in the ACTA2+ SMC module was determined using the FindMarkers function in Seurat. StringDB network analysis was performed on autosomal plaque SMC genes either higher expressed in females or males^24^. Genes were annotated with significant terms from UniProt Keywords and Gene Ontology: Biological Processes. Sex differential SMC genes and their enrichment can be found in the Supplemental Data Set.

### Data and scripts availability

Data and scripts are available upon request.

## Results

Sex-stratified patient characteristics are shown in the Supplemental Data Set. A workflow of our study and the results can be found in Fig. 1.

To determine the sex differences in gene regulation of the atherosclerotic plaque, we performed sex-differential methylation analysis in 485 carotid plaques and found differential methylation between the sexes for 4848 CpGs (sexDMPs, Supplemental Data Set). This differential methylation was present scattered over the entire autosome (Fig. 2). Mapping of the sexDMPs to genes revealed that 35.5% was located in promoters within 1kb of a transcription start site (TSS), and 23.2% was located distally intergenic. The majority of sexDMPs (86.7%) had higher methylation values in female plaques (Fig. 3A). This was replicated with 93.7% concordance in a second cohort of 190 carotid artery plaques (16.8% female) (Suppl. Fig. 1). By comparing the annotated region of all 275,204 CpGs that we tested with our 4,848 sexDMPs, we found more probes to be located distally intergenic for sexDMPs (Fig. 3B). The distribution of the location of sexDMPs was similar for those with higher methylation values in males and those in females. We allocated the sexDMPs in promoter and non-promoter DMPs and assessed their methylation state. Overall, sexDMPs showed a tendency towards a methylated state (beta-values > 0.7, Fig. 3C). However, the promoter sexDMPs were more often in a non-methylated state (beta-values < 0.3), while the non-promoter sexDMPs were more often in a methylated state. There were no differences in distribution for those sexDMPs either higher in females or males. A large proportion of sexDMPs also had methylation values between 0.3 and 0.7, highlighting that the DNA obtained from the plaques originates from multiple cells. We mapped sexDMPs to genes and performed gene enrichment analyses to determine potential underlying biology of sex differences.

Regardless of being a promoter or non-promoter sexDMP, or their methylation state, sexDMPs were enriched for development processes and cell fate commitment (Fig. 3D). We noted a large presence of transcription factors important for development, such as homeobox genes (Fig. 3E). For example, *HOXA4* contained multiple sexDMPs with higher methylation values in males in its promoter (Suppl. Fig. 2A). Another example is *SIX3*, with more than 20 sexDMPs located distally of *SIX3*, with higher methylated values in female plaques (Suppl. Fig. 2B). Even though the gene sets that we tested were enriched for similar processes, little overlap for genes was present (Fig. 3F).

### Plaque DNA methylation and cell-type deconvolution

The epigenetic landscape of DNA is cell- and tissue dependent. As plaque composition differs between the sexes, and sex differential methylation was enriched for cell fate commitment processes, we investigated cell type composition as determined by deconvolution analyses of the DNA methylation data. We dictated the algorithm to obtain 7 cell-types (see Methods). We found significant sex differences in the proportion of deconvoluted cell-types for 4 of the 7 cell-types (Fig. 4A). We correlated the proportion of deconvoluted cell-types and noted mostly negative correlations between the proportion of the different cell-types. For example, cell-type 1 was heavily negatively correlated to cell-type 6 and 7 (Fig. 4B). To determine similarities between the deconvoluted cell-types, we compared their DNAm patterns. We found strong agreement between cell-types 1 to 4, that did not correlate with celltypes 5, 6, and 7 (Fig. 4C). Cell-types 6 and 7 were again more similar to one another. We determined the potential cell-types of those with a significant sex difference. The promoter CpGs that were non-methylated in cell-type 1 and 4, but methylated in cell-type 6 and 7 were enriched for SMC-like and endothelial cell-like processes, such as muscle contraction and circulatory system process (Fig. 4D). Those promoter CpGs that were methylated in cell-type 1 and 4, and not methylated in 6 and 7, were strongly enriched for leukocyte processes and T-cell activation. Thus, we found sex differences in cell-type composition based on DNA methylation for more potential SMC-like cells in female plaques and more immune-like cells in male plaques. Plaque composition sex differences in our cohort were corroborated by more staining in female plaques for SMCs (Suppl. Fig. 3). Male plaques tend to have more presence of intraplaque haemorrhage (Suppl. Fig. 3). By laying our sexDMPs over the projected cell-types, we found that the majority of sexDMPs was similarly methylated over the cell-types (Suppl. Fig. 4). Clustering the sexDMPs over the cell-types by kMeans-clustering (k = 7) showed large differences between 2 clusters for the cell-types (Suppl. Fig. 4). All different clusters were again enriched for cell fate commitment and development processes (Suppl. Fig. 4). To determine the amount of sex differences driven by plaque composition, we reran our sex differential DNA methylation analysis corrected for cell-type proportions determined by our DNA methylation data. Instead of 4848 sexDMPs, we found 3081 sexDMPs independent of cell-type differences, of which 1649 sexDMPs overlapped (Fig. 4E). This highlighted that regardless of significant differences in projected cell-types, substantial sex differential methylation remained present on the autosome.

**Fig 4.**
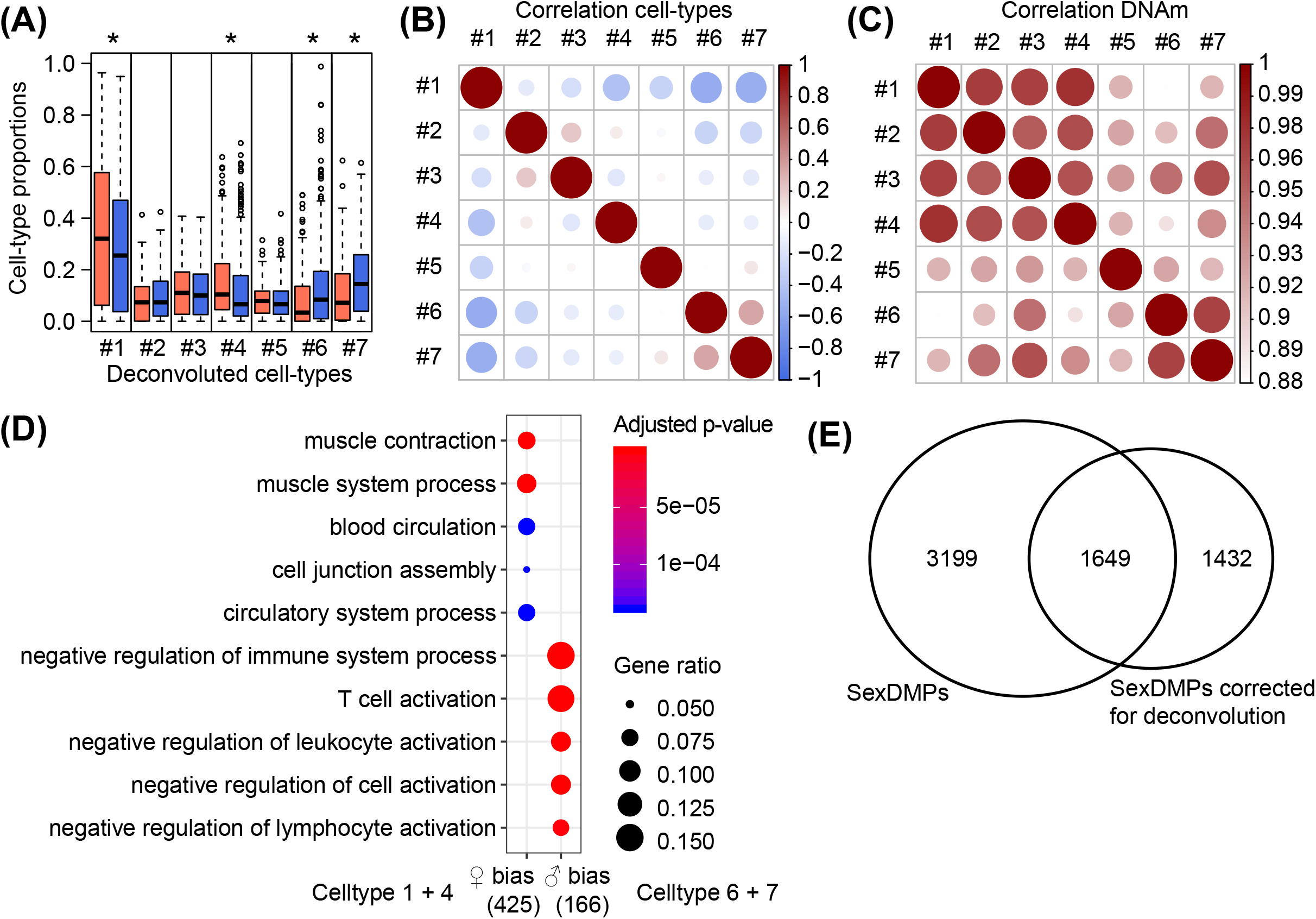
Sex differences in plaque composition by DNA methylation deconvolution. A) Barplots are shown that indicate the sex-stratified proportion of deconvoluted cell-types. An asterisk indicates a statistical significant sex difference of *p* < 0.05 (Welch Two Sample t-test). Red color indicates female plaques, blue color indicates male plaques. B) A correlation dot-plot shows the correlation coefficient between the proportion of cell-types. Color and size correspond to the correlation coefficient. C) A correlation dot-plot shows the correlation coefficient between the deconvoluted beta-values of 275204 CpGs in all cell-types. Color and size correspond to the correlation coefficient. D) A dotplot is shown for Gene Ontology enrichments for the genes mapped to by sex-biased cell-type promoter CpGs. The female bias consists of CpGs not methylated in cell type 1 and 4, but methylated in cell type 6 and 7, and vice versa for the male bias. Terms are allocated to the rows, color indicates significance, and size of the dot indicates the ratio of genes from the set present. The number indicates the number of genes that could be found in any of the sets tested. E) A venn diagram shows the overlap between sexDMPs and the sexDMPs that correspond to a sex differential DNA methylation model corrected for deconvoluted cell-type proportions.

### Sex differences in RNA expression of plaques

We performed RNA-sequencing on the same plaques to study sex differences in RNA expression and the pathology of atherosclerosis and its relation to the DNA methylation patterns of the deconvoluted cell-types. Differential expression analysis found 563 genes differentially expressed between the sexes, of which 526 were higher expressed in females (Fig. 5A, Supplemental Data Set). Higher expression of the haemoglobin genes *HBB, HBA1*, and *HBA2*, in male plaques corroborated the more frequent presence of intraplaque haemorrhage in males (Fig. 5BC). Genes that were higher expressed in female plaques were enriched for extracellular matrix organization and TGF-β signalling, terms associated with SMCs (Fig. 5D). We backtracked the female-biased genes to the DNA methylation data, and found that promoter CpGs of the female-biased genes had lower methylation values specifically in cell-type 1 and 4, the cell-types associated with SMC biology (Fig. 5E). To find out whether sex differences in RNA expression might be linked to sex differences in DNA methylation, we determined the amount of potential regulatory links for sexDMP loci. We calculated possible links based on distance and correlations between histone modifications and expression from the ENCODE Tissue Panel using T-Gene^12^. We found 29,001 potential regulatory links for 4,145 sexDMPs (correlation and distance *p* < 0.05, Supplemental Data Set). Next, we determined whether the autosomal female-biased genes (356 genes) were enriched for regulatory links based on the sexDMPs. Female-biased genes were significantly enriched in the genes targeted by the putative regulatory links (102 out of 356, hypergeometric test *p*=2.14e-8). Furthermore, we found a higher number of regulatory links for female-biased genes as compared to random genes (*p*_permutation_ = 4e-4, Fig. 5F). This indicated that regulatory regions based on sex differential methylation were associated with sex difference in RNA expression in the atherosclerotic plaque. These enrichments were different for promoter sexDMPs (*p*_permutation_ = 0.09, Suppl. Fig. 5) and distal sexDMPs (*p*_permutation_ < 1e-4, Suppl. Fig. 5), highlighting a potential differential mechanism based on distance.

**Fig 5.**
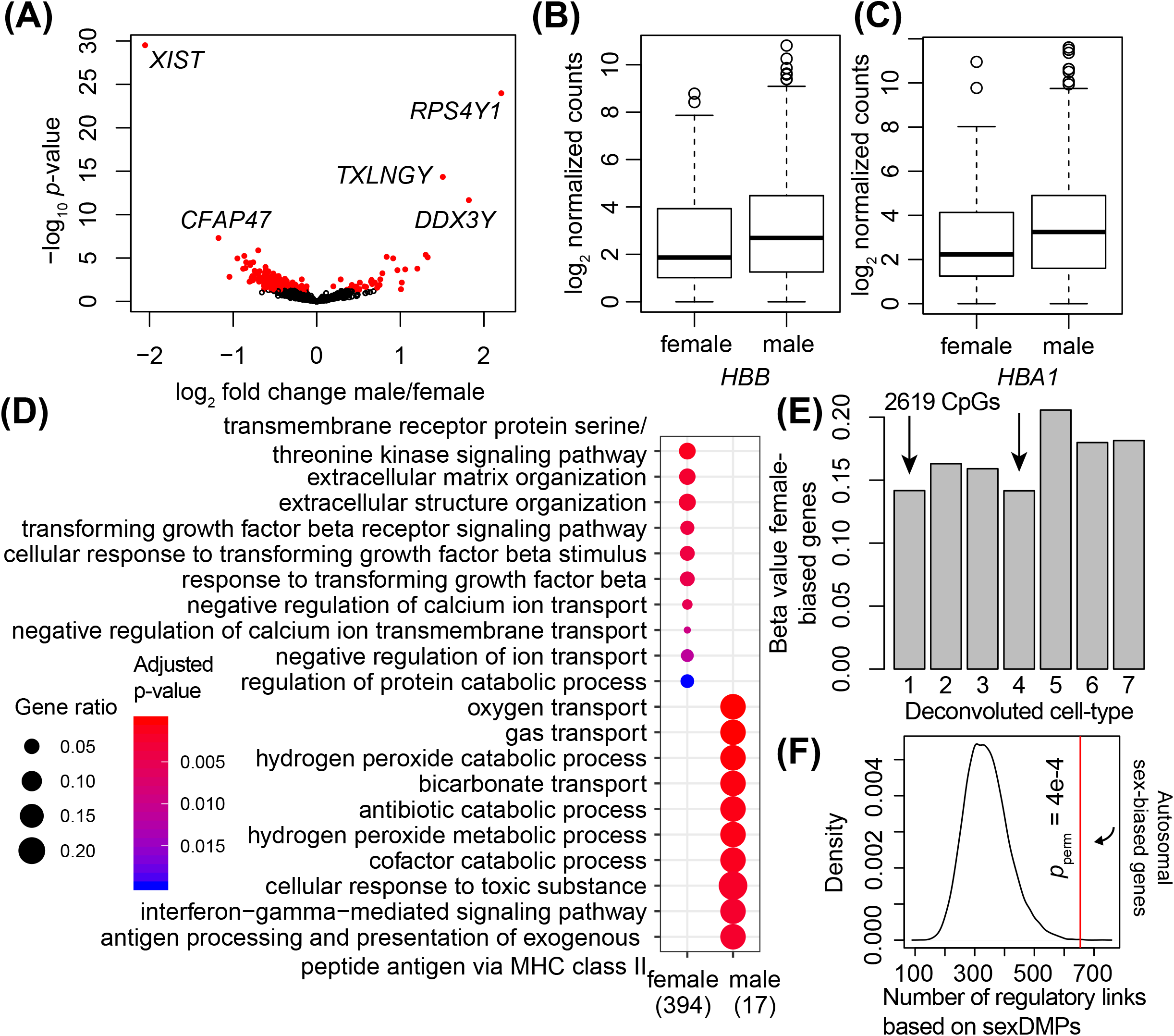
Sex differences in RNA expression of the carotid artery plaque. A) A volcano plot shows the relationship between significance and effect size by sex differential expression. X-axis indicates log_2_ fold change, y-axis indicates *p*-value. A red color highlights genes with a *p*-value < 0.05. The top sex-chromosomal genes are depicted with their gene symbol. B) Beeswarm plots show the difference in expression of the haemoglobin gene *HBB* between males and females. The y-axis shows normalized count data. C) Beeswarm plots show the difference in expression of the haemoglobin gene *HBA1* between males and females. The y-axis shows normalized count data. D) A dotplot is shown for Gene Ontology enrichments for the genes higher expressed in females (left) and the genes higher expressed in males (right). Terms are allocated to the rows, color indicates significance, and size of the dot indicates the ratio of genes from the set present. The number indicates the number of genes that could be found in any of the sets tested. E) A barplot shows the average beta-values of 2619 CpGs mapping to the promoters of genes higher expressed in females over the deconvoluted cell-types. Arrows indicate the SMC-like cell-types, highlighting that the female biased genes have lower promoter methylation in these deconvoluted cells. F) A density plot shows the permuted distribution of regulatory links with random genes. The vertical red line indicates the number of regulatory links to autosomal female-biased genes.

### Sex differences within SMCs

Sex differences in TGF-beta and extracellular matrix pathways might be driven by the differences in cell composition, but also by sex differences in SMC gene expression *per se*. To tackle this question, whether the expression differences are also present on a cellular level, we generated sex-specific single-cell RNA-sequencing data of freshly isolated atherosclerotic plaques. After expression analysis, we were able to detect 18 different cell-types. We projected the female-biased genes from the RNA-sequencing of the plaque as a conglomerated modulescore over the different plaque cell-types from the single cell RNA-sequencing. Female-biased genes were higher expressed in SMCs and endothelial cells (Fig. 6A). In addition, these genes were higher expressed in female SMCs as compared to male SMCs (Fig, 6B, *p* = 0.0003, Welch Two Sample t-test). This difference in SMC gene expression between the sexes still stands when removing the genes located on the sex-chromosomes (*p =* 0.0009). Next, we determined whether there was a significant overlap between female-biased genes and sex differential genes within the single cell SMC cluster (Supplemental Data Set). We found a significant overlap, i.e. 14 genes were overlapping (hypergeometric test *p* = 2.7e-9), indicating that at least part of sex differences in gene expression between males and females in atherosclerotic plaques is driven by sex differential gene expression in SMCs. Lastly, we performed StringDB network analysis on autosomal genes higher expressed in female plaque SMCs and male plaque SMCs (Fig. 6C). Both networks contained more interactions than randomly expected (female: PPI enrichment *p*-value < 1e-16, male: PPI enrichment *p*-value = 1.55e-15), indicating that these groups of genes are at least partially biologically connected as a group. Female plaque smooth muscle genes contained protease inhibitors, ribosomal proteins, and extracellular matrix components, whereas male plaque smooth muscle genes were enriched for responses to cytokines and lipids, as well as some extracellular matrix components (Fig. 6C & Supplemental Data Set).

**Fig 6.**
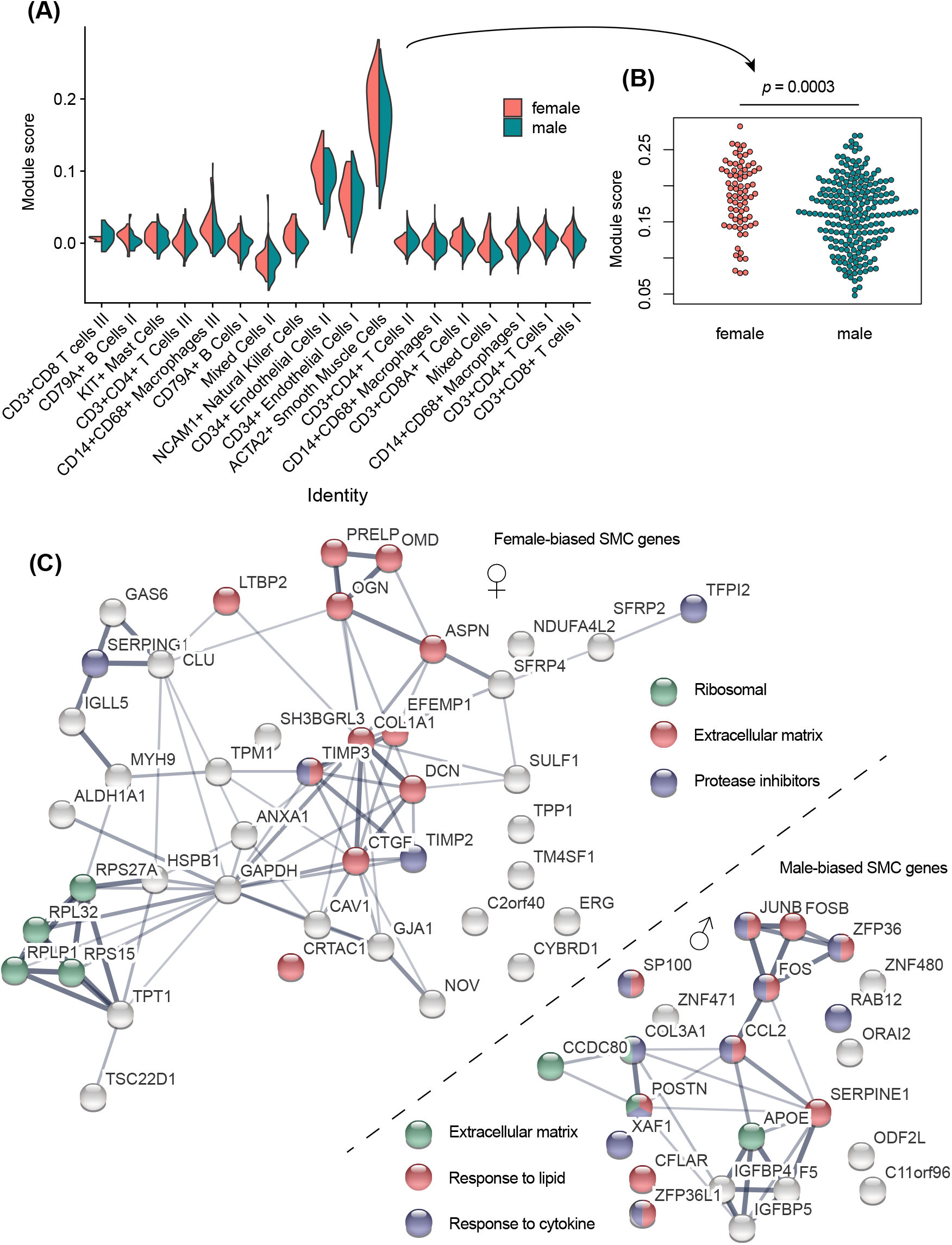
Sex differences in single-cell RNA expression of female-bias genes in the carotid artery plaque. A) A violin plot shows the conglomerated module score of the genes higher expressed in bulk plaques over 18 different single-cell RNA-sequencing derived cell-types. Pink indicates female cells, blue indicates male cells. B) A beeswarm shows the sex difference in module score in ACTA2+ SMCs (Welch Two sample T-test). C) Interactions networks based on StringDB analyses are shown for single-cell SMC-derived sex differences. Autosomal genes higher expressed in female cells are depicted in the top left, while those with higher expression in males are shown in the bottom right. Colors of the nodes show functionality of some highlighted genes based on the legend different for both sexes.

## Discussion

In this study, we performed a thorough analysis to identify sex-differential autosomal DNAm sites and the corresponding (single cell) RNA expression of carotid plaques to shed light on the understudied pathophysiology of the stable atherosclerotic plaque that is common in women. Sex differences in plaque composition based on histology and cardiac imaging have been shown before, while our knowledge on the molecular biology underlying these differences is limited. We found significant sex differences in autosomal DNA methylation and (single cell) RNA expression revealing female-specific smooth muscle biology, as well as evidence for a putative regulatory role for sex differential methylation linked to expression. We revealed that sex differential targets are mostly expressed in plaque endothelial cells and plaque SMCs, using single-cell RNA sequencing, as well as generally higher expressed in females. In addition, we found differential pathways in female- and male-biased SMC genes, that may guide future studies into the symptomatic stable plaque.

Differences in plaque rupture and plaque erosion can potentially be ascribed to differential functioning of endothelial cells and SMCs. Since plaque rupture and plaque erosion show a sex bias, it is tempting to hypothesize that sex differences in endothelial cells and SMCs underlie this bias. Using single-cell RNA sequencing, we found sex-specific SMC genes pointing to differential processes. Female-biased genes were enriched for protease inhibitors, such as *TIMP2* and *TIMP3*, proteins that can potentially halt the remodelling leading to unstable plaques, while ribosomal protein enrichment points to enhanced protein turn-over. Male smooth muscle genes were enriched for immediate early response transcription factors, such as *JUNB* and *FOSB*, as well as responses to lipids and cytokines, highlighting that male and female plaque SMCs may present with sex-specific biology. Extracellular matrix proteins constitute the majority of proteins within the plaque, such as OGN, ASPN and DCN^25^. We report here that these genes show sex difference in expression in SMCs within the plaque. Previous efforts have elucidated extracellular matrix proteins that discriminate between symptomatic versus asymptomatic lesions based on plaque rupture as well^26^. Some of the most differential proteins in these studies (encoded by genes *COL3A1, TIMP2, TIMP3, ASPN*) are sex-differential in our analyses, underscoring that these genes might be involved in sex differences in plaque stability. Of note, we have recently seen that atherosclerotic plaques are becoming more stable, i.e. plaques show a reduction in paradigmatic measures of plaque vulnerability^27^. However, the biology of the symptomatic stable plaque is not as well understood as the symptomatic unstable plaque^28^. Since women suffer more from stable plaques and erosion, research efforts into sex differences will be of benefit to the future treatment of atherosclerosis.

Our data not only identify potential sex-specific relevant genes for the atherosclerotic disease trajectory and their cell-type, but also underscore the importance of looking at sex as a biological variable explaining different phenotypes underlying atherosclerotic disease. However, sex is not often taken into consideration as an outcome in cardiovascular epigenetics, even though epigenetic phenomena are influenced by sex, as we have highlighted here as well.

Sex differences in DNA methylation may lead to sex differences in gene regulation. DNA methylation is known to affect gene regulation, and it is commonly accepted that methylated promoters are a hallmark of gene repression. However, there are also studies that suggest that methylation correlates with gene expression^29^. More and more transcription factors are being discovered that can bind methylated DNA, such as Kaiso and KLF4, and may pioneer gene expression^30^. Sex differences in DNA methylation might lead to differences in transcription factor binding and subsequent gene regulation. This is underlined by the fact that regulatory links based on sexDMPs are enriched for genes higher expressed in the female plaque as compared to the male plaque.

Irrespective of plaque composition differences, DNAm differences in the autosome may underlie the sex differences in the development of atherosclerosis. Actually, the majority of CpGs (86.7%) with a sex difference in methylation, had higher methylation values in females. This cannot be explained by the normalization procedure we performed, as normalization is performed based on probes designed for normalization. Mamrut et al. also found the bulk of the significant CpGs to be hyper-methylated in females as compared to males^31^, which agrees with our finding. Enzymes that maintain DNA methylation might be more active in females, be present in more numbers around the autosome, or are affected by proteins and molecules present in different levels between the sexes, such as sex hormones and their receptors, and sex chromosomal gene products. One might speculate that these differences are driven by sex differences in chromatin conformation, because of the differential action and three-dimensional spacing of the sex chromosomes. However, knowledge regarding sex differences in complete chromosome conformation is lacking.

DNA methylation is essential for maintaining genome stability. Large differences in methylation of the epigenome, with the bulk being hypermethylated in females, indicate that male cells might be more prone to genomic instability than female cells. This may perhaps partly explain why men develop non-reproductive cancers earlier than females^32^. Males also develop coronary artery disease 7 to 10 years earlier than females, which is preceded by the formation of atherosclerotic plaques, a process that has a lot in common with cancer^33^. Sex differences in genomic instability caused by differences in (the metabolism of) DNA methylation might affect sex-bias in atherosclerotic disease.

To see how our results compare to previous studies on sex differences and DNA methylation, we overlapped sexDMPs with previously found CpGs significantly affected by sex. The study of Singmann et al.^34^ looked at whole-genome autosomal DNAm differences between the sexes in blood, and found 1,182 significant CpGs after meta-analysis. Of the 1,182 CpGs, 13 overlapped with the 4,848 sexDMPs. Another study by Mamrut et al.^31^ investigated DNAm differences between the sexes in blood as well, and found 7,454 significantly different autosomal CpGs. Of these 7,454, 55 overlapped with our 4,848 sexDMPs. Zero CpGs of the 13 and 55 overlapping CpGs were shared.

The minor overlap between the two other studies and the present study points to a low number of CpGs consistently differentially methylated between the sexes over different cell-types, as the two other studies measured DNAm in blood, and perhaps as well to sex differences in different health or disease states.

### Sex hormones and DNAm sex differences in atherosclerosis

An explanation for the sex differences found in these epigenetic markers might be sex hormones. Our population is of older age, with a mean age for both men and women around 68 years of age (Supplemental Data Set). This indicates that most of the women within our cohorts have been post-menopausal for years. Estrogen levels between the men and women in our cohort do not differ (Supplemental Data Set and ^35^). However, estrogen could have affected the methylation state of some of the sex-CpGs found, as an entire reproductive life cycle of fluctuating estrogen levels is bound to change the epigenetic landscape of cells. Whether or not these changes are still in effect years after menopause is not known, but is has been suggested that menopause *per se* accelerates epigenetic aging^36^. Furthermore, sex hormones and sex hormonal receptor action affect plaque composition^37–39^, which will inadvertently affect sex differences in DNA methylation. However, sex hormones have been shown to affect different epigenetic markers and sex hormone receptors have been shown to bind different epigenetic regulators^40,41^. Testosterone levels do differ between the sexes and it has been shown that testosterone has an effect on plaque characteristics within males^35^. Unfortunately, we could not correct our results for testosterone levels, as these were in range of the detection limit of the assay for women, and thereby prone to erroneous interpretation.

### Strengths and limitations

The Athero-Express Biobank is the largest plaque biobank worldwide which allowed us to investigate sex differences with sufficient numbers of women. However, a limitation in our study is that our cohort contains more plaques from men than from women. The biobank contains advanced atherosclerotic plaques, limiting us to describe the sex differences found in the end-stage culprit lesions, not properly reflecting the development of the atherosclerotic plaque. Targets for initiation and progression of atherosclerotic disease may differ between the sexes, which unfortunately we cannot address. Additionally, we realize that the sex differences we describe might at the same time also be subject to gender differences, something we could not address in the current study.

## Conclusion

We conclude that there are profound sex differences in DNA methylation and RNA expression in human atherosclerotic plaques from patients undergoing carotid endarterectomy. The sex differential genes we found are prime targets for future studies into the symptomatic stable lesion that are common in women. Our evidence suggests that smooth muscle cells in the plaque are subject to these differences, as underlined by deconvolution analyses, regulatory linkage, RNA-sequencing and single-cell RNA-sequencing data. Lastly, our findings highlight the importance of considering sex as a biological variable in cardiovascular disease and genomics.

## Abbreviations

CVD: CVD Cardiovascular disease
CpG: CpG-dinucleotide
DNAm: DNA methylation
SexDMP: Sex-differentially methylated CpG-Probe
SMC: Smooth muscle cell

## Acknowledgements

We thank Utrecht Sequencing Facility for providing sequencing service and data. Utrecht Sequencing Facility is subsidized by the University Medical Center Utrecht, Hubrecht Institute, Utrecht University and The Netherlands X-omics Initiative (NWO project 184.034.019). We acknowledge the service of Single Cell Discoveries for single-cell RNA-sequencing of human plaque material.

