## Supplementary material for "Sex-dependent gene regulation of human atherosclerotic plaques by DNA methylation and transcriptome integration points to smooth muscle cell involvement in women": Suppl. Fig.

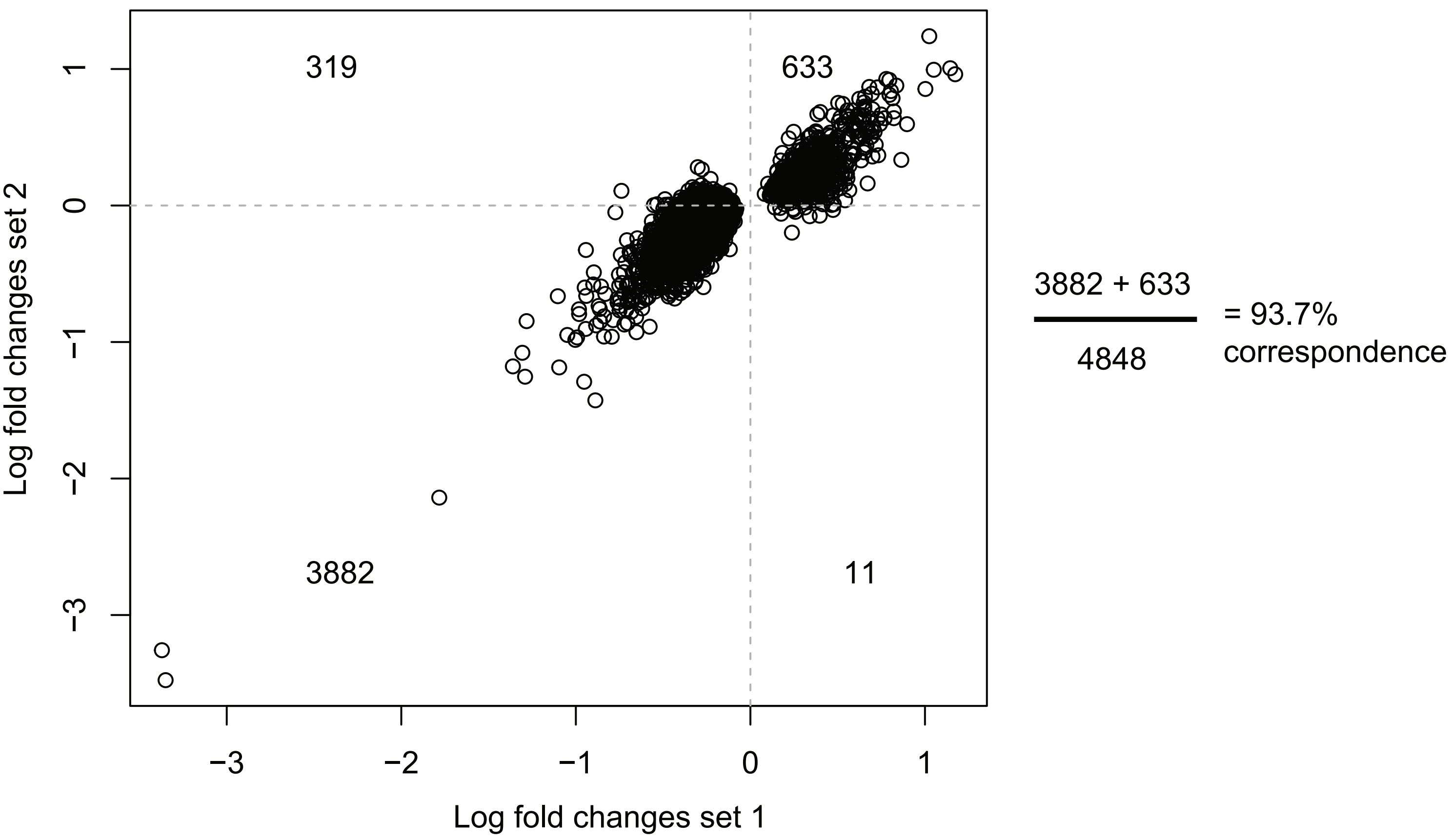


**Suppl. Fig. 1. Replication of sex differential methylation.** A scatter-plot shows the concordance between the effect size from set 1 (x-axis, 485 patients, 30.5% female) and set 2 (y-axis, 190 patients 16.8% female). The number in each quadrant indicates the dots in each quadrant.

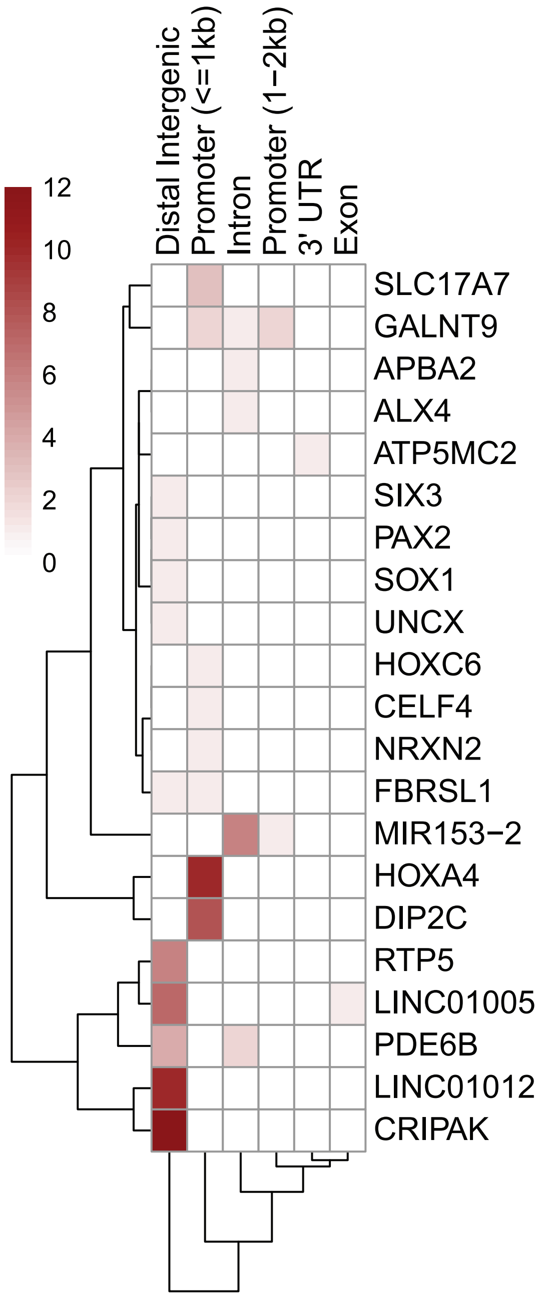


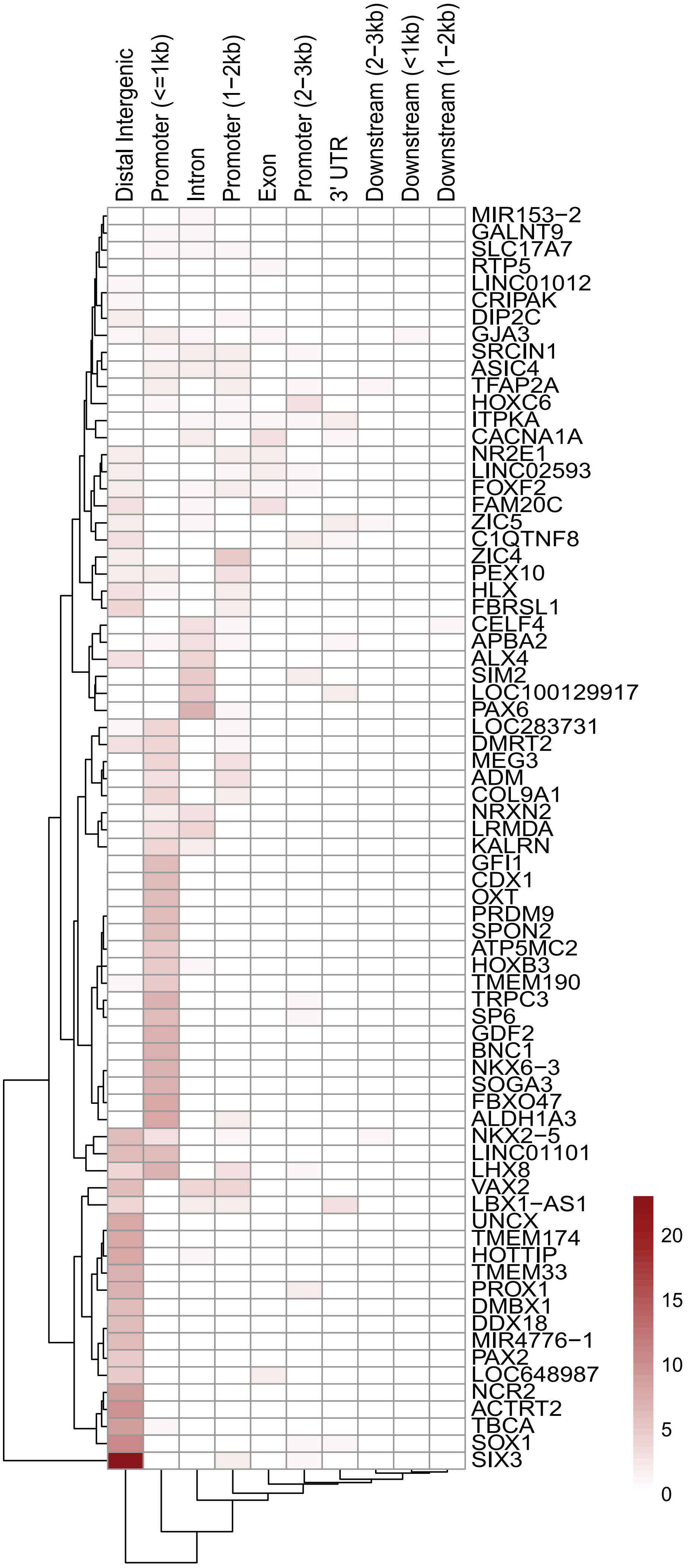


**Suppl. Fig. 2. Top genes sexDMPs.**
A heatmap shows the number of genes mapped to by sexDMPs more methylated in males (top) and in females (right) with genes on the rows and the location on the columns. Only genes that are at least mapped to by at least 5 sexDMPs are shown.

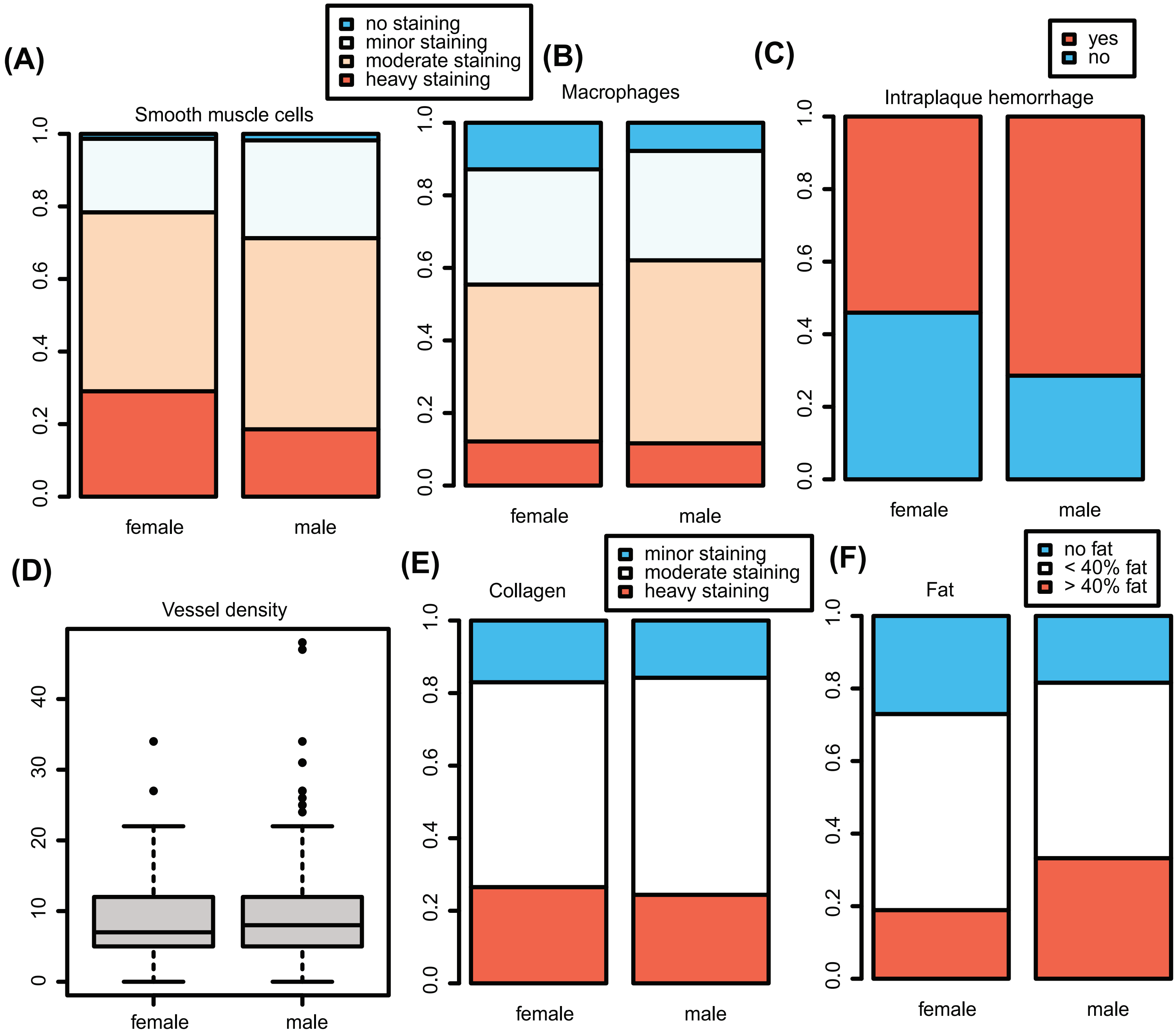


**Suppl. Fig. 3. Sex differences in plaque characteristics.** A-F show smooth muscle cell content, macrophage staining, presence of intraplaque hemorrhage, vessel density, collagen staining and fat percentage, respectively, in female (left of each panel) and male (right of each panel) samples.

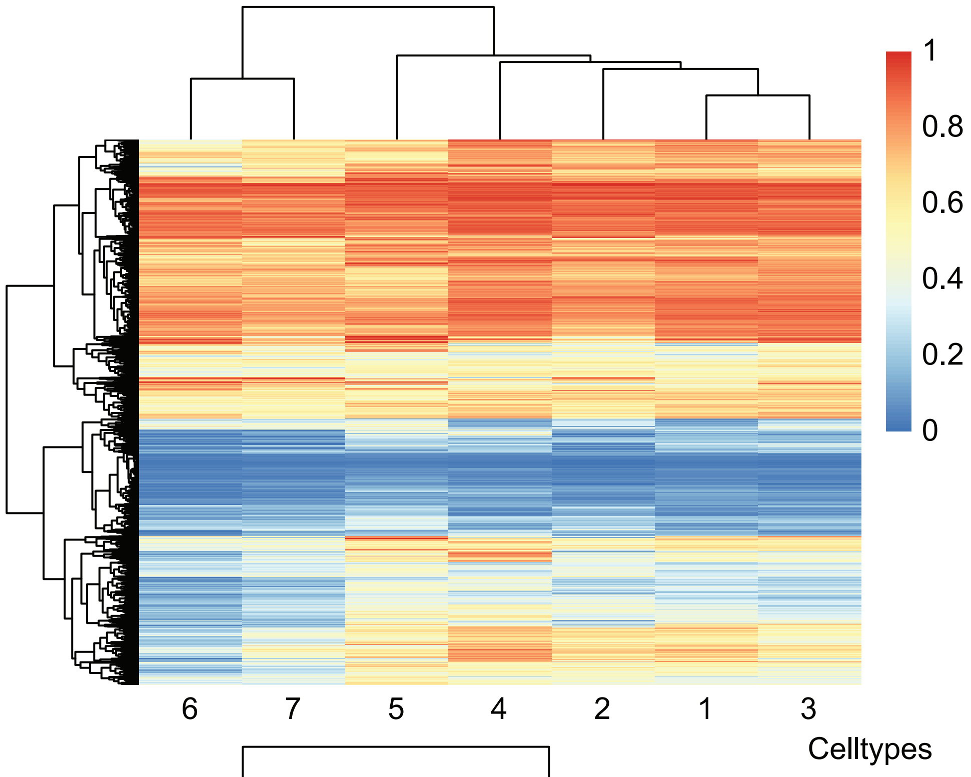


**Suppl. Fig. 4. Sex DMPs and celltypes.** A) A heatmap shows the DNA methylation pattern of the sexDMPs over the deconvoluted cell-types. The rows contain the sexDMPs, the columns show the cell-type. B) The same heatmap as A) is shown, but this one is clustered by k-means clustering (k = 7). C) A dotplot is shown for Gene Ontology enrichments for the k-means sexDMP clusters. Terms are allocated to the rows, color indicates significance, and size of the dot indicates the ratio of genes from the set present. The number indicates the number of genes that could be found in any of the sets tested.

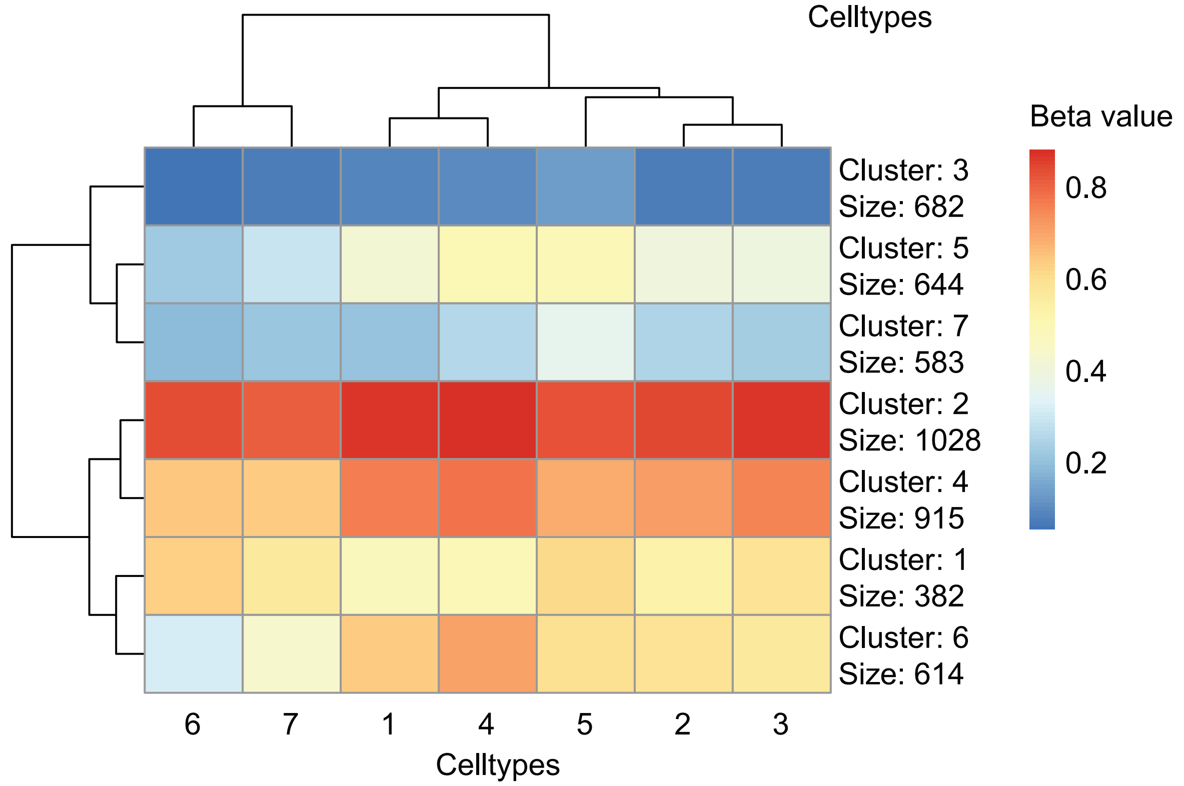


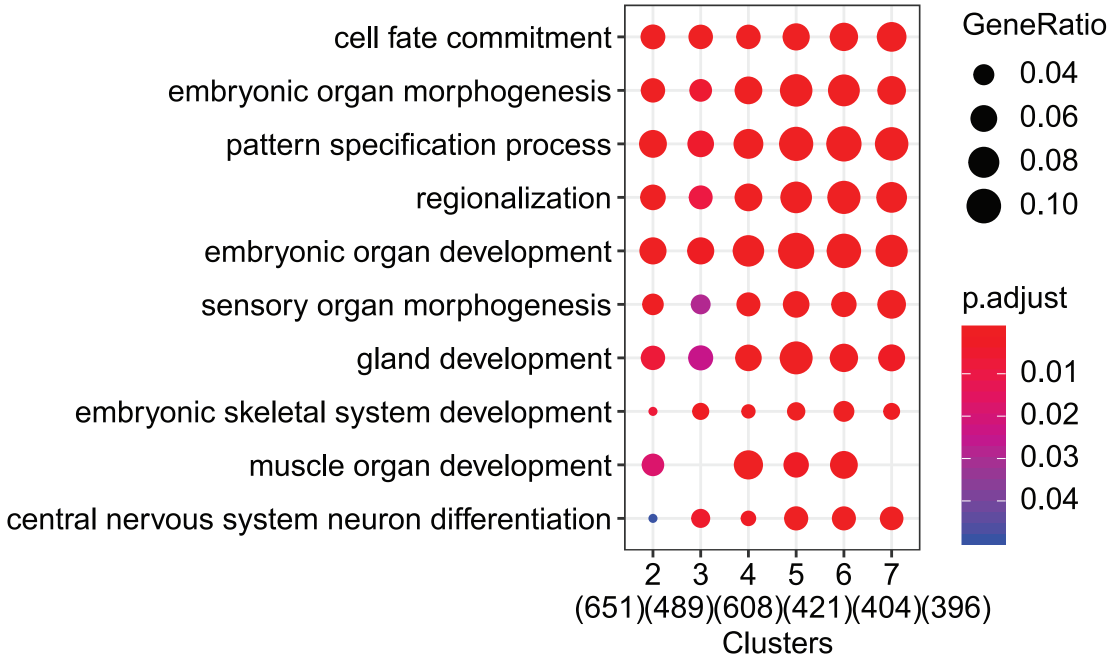

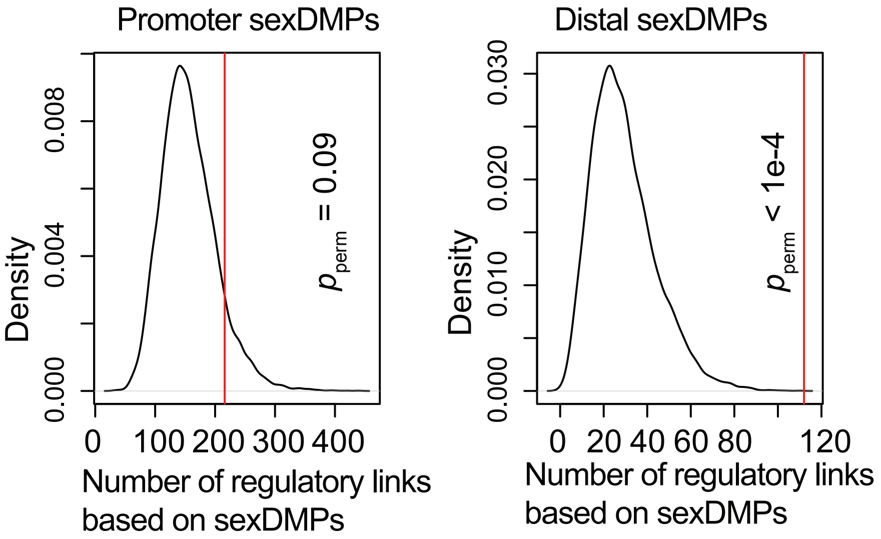


**Suppl. Fig. 5. Regulatory loci and location of sexDMPs.**

A density plot shows the permuted distribution of regulatory links based on promoter sexDMPs (left) and distal intergenic sexDMPs (right) with random genes.

The vertical red line indicates the number of regulatory links to autosomal female-biased genes.
